# ProRafts: A machine-learning predictor for *raftophilicity*, the protein affinity for biomembrane rafts

**DOI:** 10.1101/2023.03.20.533471

**Authors:** Deniz Yurtsever, Can Keşmir, Maria Maddalena Sperotto

## Abstract

**Background:** Protein raftophilicity refers to the affinity of proteins for cell biomembrane lipid domains, called ‘rafts’. Rafts are fluctuating nanoscale platforms that are enriched in cholesterol and sphingolipids, and that are considered relevant for cell signalling, viral function, and biomembrane trafficking. The dynamic partitioning of proteins into rafts depends on the physical and physico-chemical properties of the biomembranes where such proteins are embedded or attached; however it also depends on specific protein “features”, such as acylation, glypidation, specific amino acid sequence motifs, transmembrane hydrophobic length, and surface accessible area to solvent. In this paper we present a method, and the resulting “ProRafts” predictor, that can be used to predict if a given mammal protein may be “raftophilic” or “non-raftophilic”, without having an a priori knowledge of the physical and physico-chemical properties of the biomembranes where such protein is embedded or attached. ProRafts is based on a machine-learning algorithm, XGBoost, where data regarding the features of known raftophilic human-proteins fed the algorithm.

**Results:** ProRafts enabled to predict correctly more than 80% of human proteins that are *a priori* known to be raftophilic; this is a promising result considering the limited size of the training dataset that we could build with data retrieved from protein databases. In addition, although we used protein features of known human raftophilic proteins, it was possible to identify accurately raft-proteins from other *mammals* than humans, such as mouse and rats. This finding suggests that certain protein features are sufficient to predict raftophilicity of proteins from different species. Moreover, our results indicated that phosphorylation may play a more relevant role for protein raftophilicity than indicated by previous studies.

**Conclusion:** Raftophilic proteins can be used as biomarkers in medical research, or can serve as targeting sites for therapeutics. In this respect, the machine learning method presented in this paper is a useful tool to guide experimental validations of raftophilicity of proteins in biomembranes, and facilitate the choice of proteins that can be used for experiments on biomimetic membranes.

## INTRODUCTION

In the 70’, in a paper that described a revised version of the so-called Fluid-Mosaic Membrane Model (Singer and Nicolson, 1972), it was hypothesized that inhomogeneous multicomponent domain-like structures were present in biomembranes (Nicolson, 1976). Around the same time, formalized domain-theories were proposed (Stier and Sackmann, 1973; Estep et al., 1979; Karnovsky et al., 1982; Israelachvili et al., 1980), and domain-like structures were detected both in reconstituted model membranes (Estep et al., 1979) and in cell membranes (Goodsaid-Zalduondo et al., 1982). Different types of domains were identified, ranging from curved caveolae and clathrin coated pits, to flat structures enriched in specific bio-molecular components (Semrau and Schmidt, 2009). In 1997, Simons and Ikonen adopted the term ‘rafts’ to indicate those biomembranes domains that appeared to be resistant, at low temperature, to Triton X-100 detergency—hence the name detergent-resistant membranes, in contrast to detergent-soluble membranes. Because these domains were enriched in cholesterol and sphingolipids, it was suggested that they were in the so-called liquid-ordered (*lo*) phase (Ipsen et al, 1987; Quinn and Wolf, 2009; Sodt et al., 2014), and floated (like rafts in water) within the liquid-disordered, namely fluid plasma membranes—where the term “liquid” refers to the acyl chains conformation and the term “ordered”/”disordered” to the lateral membrane organization. In 2006, during a Keystone Symposium of Lipid Rafts and Cell Function, rafts were then formally acknowledged and described as “*small (10-200nm), heterogeneous, highly dynamic, sterol- and sphingolipid-enriched domains that compartmentalize cellular processes. Small rafts can sometimes be stabilized to form larger platforms through protein-protein interactions*” (Pike 2006). However, since formally acknowledged, rafts’ existence was questioned and a number of discussions and controversial issues (Magee and Parmryd, 2003; Jacobson et al. 2007; Kenworthy 2008; Nicolson, 2014; Tanner et al., 2011, Simons and Gerl, 2010) were raised, not least those related to the choice of techniques adopted to detect rafts’ presence in biomembranes (Foster and Chan, 2007). Only in the case of caveolae there was/is a general consensus that these are raft-like structures enriched in sphingomyelin, cholesterol, and caveolin protein (Parton and Simons, 2007); caveolae are relatively large (30-50 nm) and stable structures and can be isolated without using detergents (Westermann et al., 1999), thus facilitating their experimental detection. In 2010, Simons and Gerl took into account the criticism regarding how rafts-related proteins are detected, and presented what they called “an evolving model” for membrane rafts. According to this model rafts are “*defined as dynamic, nanoscale, sterol–sphingolipid-enriched, ordered assemblies of proteins and lipids, in which the metastable raft resting state can be stimulated to coalesce into larger, more stable raft domains by specific lipid–lipid, protein–lipid and protein–protein oligomerizing interaction*” (Simons and Gerl, 2010) In a subsequent paper, Simons and Sampaio (2011) used the term *raftophilicity* to indicate the affinity of both peripheral- and membrane proteins for rafts. It is worth noticing that after four decades following the publication of the Fluid Mosaic Model and its revised version (Singer and Nicolson, 1972; Nicolson, 1976), one of the authors (Nicolson, 2014) stressed again the importance of the nano- and submicro-sized domains—such as lipid rafts and membrane-associated cytoskeletal and extracellular structures—for maintaining specialized membrane structures in biomembranes, thus constituting suitable sites for bioprocesses.

Although rafts have been detected experimentally in biomimetic membrane, i.e., synthetic membranes made of lipids and proteins known to be associated with rafts (Bathia et al., 2016), advancements in biochemical and biophysical technologies, ranging from spectroscopy to microscopy, enabled to gain insight into the role of rafts in vivo, as well (Carquin et al., 2016; Sezgin et al., 2017; Owen et al., 2016, Levental, 2020; Levental et al., 2020). In particular, it was found that rafts can affect cell migration (Schug et al., 2012) and endocytosis, and can play a role in cellular signalling (Douglass and Vale, 2005), as reviewed by Cheng and Smith (2019), and subcellular trafficking (Diaz-Rohrer et al., 2014a). Rafts seem to play a role, too, in both binding, endocytosis, and replication of viruses, such as influenza, SARS-CoV2, and HIV viruses to host cells (Verma et al., 2018; Guo et al., 2017; Baglivo et al., 2020; Giese et al., 2006; Sengupta et al., 2019; Yang et al., 2017; Lajoie and Nabi, 2007; Wei et al., 2020, Lorizate and Kräusslich, 2011). A plethora of studies indicated that rafts are also implicated in processes related to aging and neurodegeneration (Marin et al., 2013; Schengrund, 2010; Antoninia et al., 2017, Colin et al., 2016; Korade and Kenworthy, 2008; Cartucci et al., 2017; Fantini and Barrantes, 2009; Santos et al., 2016). The fact that a number of membrane signalling proteins, which are involved in different types of neuropathologies, interact with lipid rafts (see for example Table 1 in Marin et al., 2013), brought some authors to suggest that rafts can be viewed as “strategic observation posts” for pathophysiological events of neurodegenerative diseases (Marin et al., 2013), and that aging and gender may cause “lipid raft aging” (Diaz et al, 2018). As rafts act as signalling and protein platforms, they may be used, too, for identifying biomarkers that fluctuate in disease cell states. To this end, rafts could be used as targeting sites for anti-cancer therapeutics – as described in the review by Staubach and Hanisch (2011). In addition, in a review Hryniewicz-Jankowska et al. (2014) discussed the dependence of several growth factor receptor signalling pathways on membrane rafts, as well as the involvement of rafts with regard to the invasiveness of cancer cells and formation of metastasis. Because of their raft-disrupting property, known synthetic and naturally occurring agents, such as filipins and statins, are listed as potential anticancer therapeutics.

**Table 1.** **General protein features as** retrieved from/by: ^**1**^ UniProt DB, ^**2**^ modlAMP package, and ^**3**^ Biopython package – ProtParam.

| General and sequence-motif Protein Features | Definition |
| --- | --- |
| Protein sequence length<sup>1</sup> | The number of amino acids in the canonical sequence. |
| Molecular mass<sup>1</sup> | The mass of the protein in Daltons (Da). |
| Aliphatic index<sup>2</sup> | The relative volume of the protein occupied by aliphatic side chains (Alanine, Valine, Isoleucine, and Leucine). |
| Net charge (pH 7.4)<sup>2</sup> | The total molecular charge at pH 7.4. |
| Boman index<sup>2</sup> | The estimation of potential protein-membrane interaction. |
| Isoelectric point<sup>3</sup> | The pH at which the protein carries no net electrical charge. |
| Amino acid composition<sup>3</sup> | Fraction of amino acids present in the sequence. |
| Hydrophobicity<sup>3</sup> | The sum of amino acid hydropathy values. |
| Number of +/- amino acids<sup>3</sup> | The total number of positive and negative amino acids. |
| CRAC, CARC motifs<sup>3</sup> | Cholesterol recognition/interaction amino acid consensus patterns. |

### Rafts proteomics

Already in the early 80’s Karnovsky and co. (1982) asked themselves the question: “Do specific membrane proteins reside in specific lipid domains, and can perturbation of the specific domains structure affect protein structure and function?”. However, and as Rabilloud (2009) poetically stated, almost three decades later, when it comes to membrane proteins and proteomics, one should keep in mind that ‘Love is possible, but so difficult’. This difficulty arises from the fact that membrane proteins requires both a hydrophilic and a hydrophobic environment in order to perform biological functions, but protein chemistry methods work mainly in water-based media. Therefore, alternative methods are needed and, as suggested by Diaz (2010), computational and bioinformatics tools may be used to investigate protein raftophilicity. To take advantage of such tools—without having an *a priori* knowledge of the physical and physico-chemical properties of the membranes and the domains where the protein may be embedded or attached —one could start by identifying protein “features” that facilitate targeting to rafts. These features may refer to physical, and structural properties, post translational modifications, cellular localization, and sequence and sequence-related properties.

### Protein features that may facilitate raftophilicity

As was illustrated by Pike in 2004, and shown in Figure 1, proteins partitioning into lipid rafts may depend on proteins lipidation, glypiation, transmembrane domains length, and the interaction of specific amino acid sequence motifs with the lipid components of rafts such as cholesterol (Epand et al., 2010; Simons and Sampaio, 2011) and sphingolipids (Fantini, 2003; Mahfoud et al., 2002). In particular, there are strong indications that proteins’ post-translational modifications by saturated fatty acids facilitate raftophilicity (Levental et al., 2010a,b; Sezgin et al., 2017); however, farnesylation and geranylgeranylation, namely modifications by short, unsaturated, and/or branched hydrocarbon chains, do not favour raftophilicity (Melkonian et al., 1999). GPI-anchors direct proteins to lipid rafts using the two fatty acids within the GPI group as anchor (Lorent and Levental, 2015; Lorent et al., 2017);

**Figure 1.**
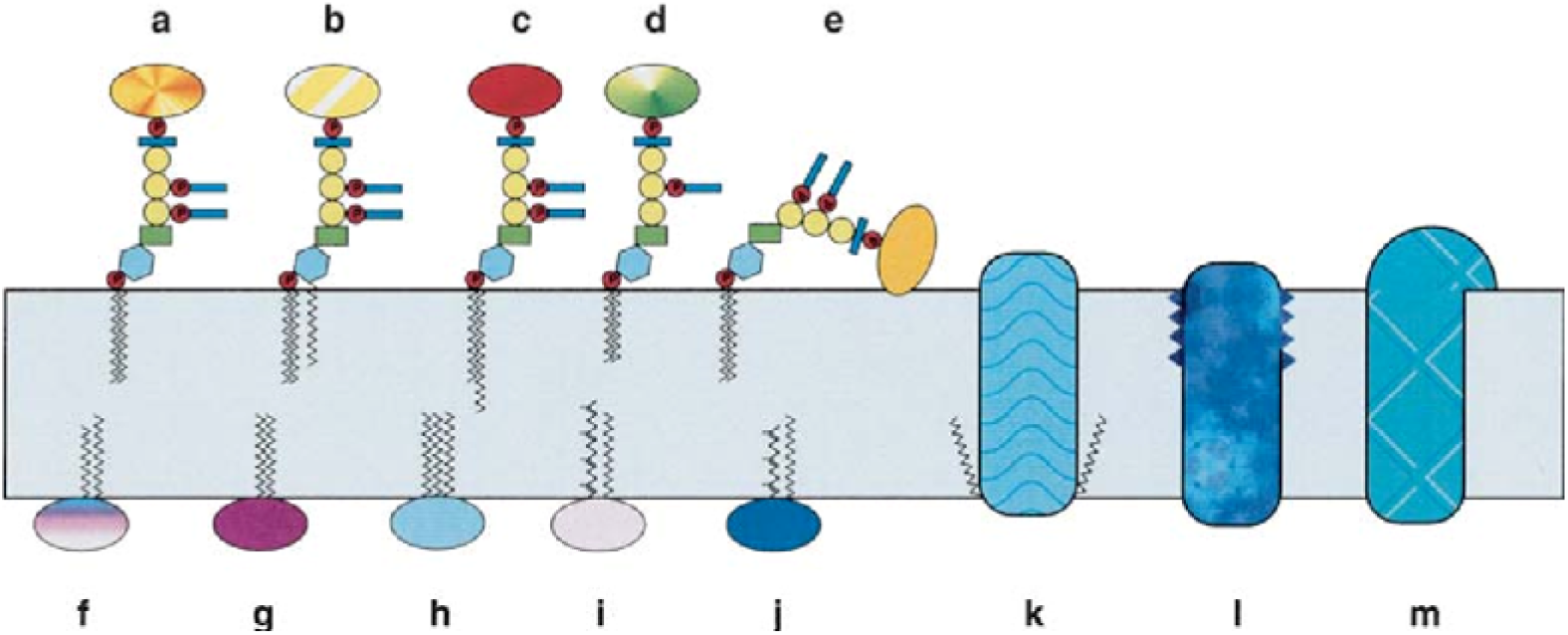
Protein types that may target rafts: **a-e**, GPI-anchored proteins. **f**, proteins modified by attachment of myristoyl group and palmitic acid. **g-h**, proteins modified by attachment of two and three palmitic acids, respectively. **i**, proteins modified by attachment of a geranylgeranyl group and palmitic acid. **j**, proteins modified by attachment of farnesyl group and palmitic acid. **k-m**, transmembrane proteins that interact with rafts via (**k)** two attached palmitic acids, and (**l, m)** amino acid residues. Figure from “Lipid rafts: heterogeneity on the high seas” by L.J. Pike, 2004, *Biochemical Journal, 378*(2), 281-292.

Proteins may also be targeted to rafts because of their direct interaction with their lipid components. As cholesterol is enriched in rafts, and plays a role, too, in many membrane-associated processes where proteins are involved, attempts were made to find cholesterol-recognition amino-acid sequence motifs in both peripheral- and membrane proteins (Di Scala et al., 2017).The existence of a common cholesterol recognition/interaction amino acid consensus pattern–acronym CRAC–was first postulated by Li and Papadopoulos (1998) who investigated the functioning of the peripheral-type benzodiazepine receptor, an outer-mitochondrial membrane protein involved in the transport of cholesterol from the outer to the inner mitochondrial membrane. The CRAC motif, which exhibits a preference for the inner leaflet of membranes, is of the type (L/V)-X_1–5_-(Y)-X_1–5_-(K/R), where the neutral and hydrophobic L or V are the branched apolar amino-acid residues, X indicates any residue (1 to 5 sequence length), Y is a mandatory aromatic residue position such as tyrosine, and a basic amino acid such as Arginine or Lysine. Although a protein containing the CRAC-motif is not always targeted to rafts (as the motif is also found in prokaryotes, which are depleted of cholesterol), the CRAC motif is nevertheless abundant among proteins that interact with cholesterol (Epand, 2006; Epand et al., 2010), such as gp41 (the fusion protein of HIV) and caveolin. In membrane proteins, another cholesterol-recognition motif was also detected (Baier et al., 2011): the CARC sequence motif, (K/R)-X_1–5_-(Y/F)-X_1–5_-(L/V). This motif constitutes an inverted CRAC, as the sequence is similar to the one of CRAC but oriented in the opposite direction along the polypeptide chain. Compared to the CRAC motif, which exhibits a preference for the inner membrane leaflet, the CARC exhibits a preference for the outer leaflet (Fantini et al., 2016). Both motifs are characterized by a positively charged basic residue at one terminal and an apolar amino acid residue at the other end (Fantini and Barrantes, 2013). Recent studies by Listowski et al. (2015) also showed that the CRAC and CRAC-like motifs, which are found in the surface of the human Membrane Palmitoylated Protein 1 (MPP1)—which is a major target of palmitoylation in red blood cells—are responsible for binding cholesterol, and thus may be implicated in rafts. It was also suggested that protein hydrophobic length may play a role for protein affinity for raft, at least for the case of multi-transmembrane strand proteins (Lin and London, 2013). This is because, to minimize the hydrophobic mismatch between the lipid bilayer thickness and the protein hydrophobic length, proteins with longer transmembrane domains (TMDs) preferentially partition into the thicker biomembrane rafts. Furthermore, a recent study (Lorent et al, 2017), performed on dozens protein TMD of single-pass proteins, confirmed that, besides protein hydrophobic length, two other protein features, such as the transmembrane domain surface accessible area (ASA)—where proteins with smaller ASA prefer to partition into rafts than those with larger ASA—and palmitoylation, may influence raftophilicity. Experiments by the same authors showed that Post-translational Modifications (PTMs) located closer to the TMD have larger influence on raftophilicity than PTMs located further away. Based on the observation that the same protein features that determine raftophilicity were also found to determine plasma membrane localization (Sharpe et al., 2010), Lorent et al. (2017) proposed a mechanist explanation—supported by experimental observations (Sharpe et al., 2010; Munro, 1995)—for plasma membrane localization being driven by raft-mediated sub-cellular sorting/trafficking, as previously also hypothesized by Diaz-Rohrer et al. (2014a, 2014b).

### Bioinformatics and protein raftophilicity

In 2002, Jensen et al. developed an Artificial Neural Network (ANN) method that identifies and integrates relevant features that can be used to assign proteins of unknown function to functional classes, and enzyme categories for enzymes. The idea was to use a number of proteins’ functional attributes together with proteins’ amino acid sequence for training an ANN, hence make functional predictions on unknown protein sequences. The work presented in this paper was inspired by this type of feature-approach. We developed a mammalian-protein raftophilicity prediction method, which makes use of a machine learning algorithm, the eXtreme Gradient Boosting, in short XGBoost (Chen and Guestrin, 2016). The outcome is a predictor, ProRafts, which can classify human (and other mammals) proteins into one of the two categories, (raftophilic, non-raftophilic), and can identify the protein features that may contribute significantly to raftophilicity.

## METHODS

Figure 2 shows the flowchart for the training, testing and validation process used during the development of the ProRafts predictor. For training, testing, and validation of the predictor we used both a positive (raftophilic) and a negative (non-raftophilic) dataset. The training dataset was used to train the XGBoost algorithm, with hyper-parameter tuning, and the validation dataset was used for validation. If the validation results were not satisfactory, the predictor was re-trained via hyper-parameter tuning, and then the predictor was tested on the independent testing dataset.

**Figure 2.**
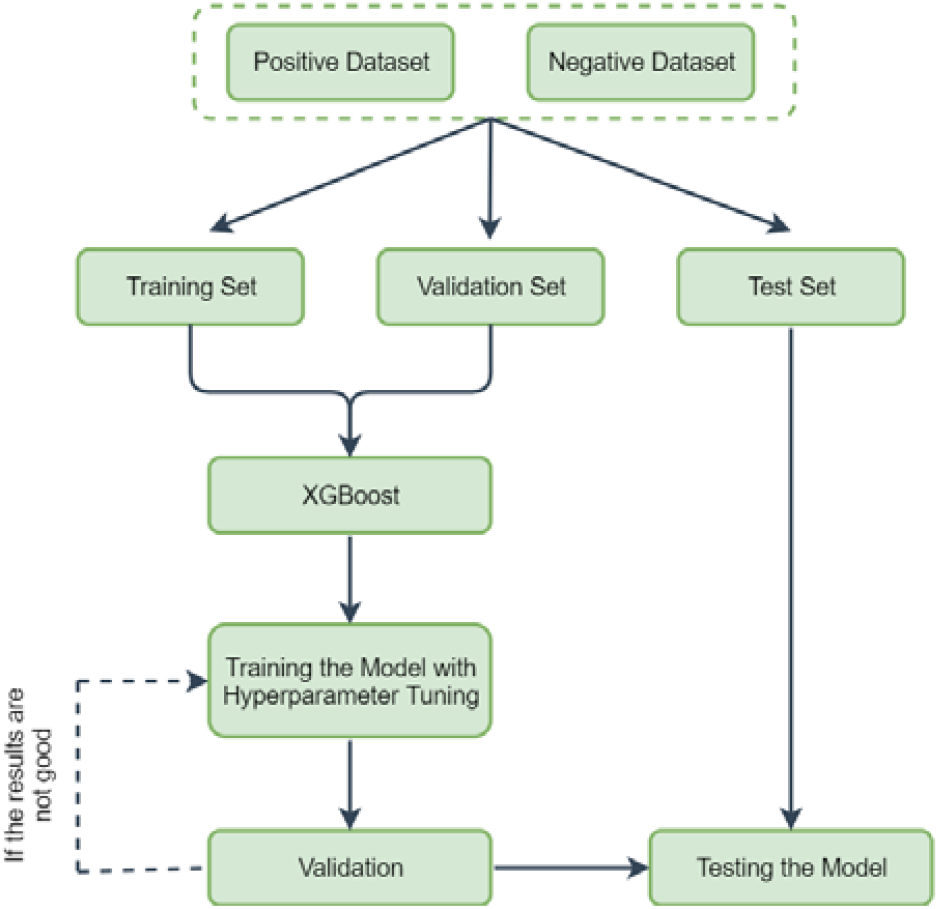
Flowchart of the machine learning training process. Both a positive dataset and a negative dataset were used for the training, validation and testing the model, *i*.*e*., the predictor. The training dataset was used to train the predictor with hyper-parameter tuning, and the validation dataset was used for validation. When the validation results were not satisfactory, the predictor was re-trained with hyper-parameter tuning and then it was tested with an independent test dataset.

### Retrieval of positive and negative datasets

The data of the positive dataset for training and testing the predictor were retrieved from RaftProt V2 (Mohamed et al., 2019). RaftProt V2 is a protein database from published studies regarding mammalian lipid raft proteomics. The reported proteomics data originates from mainly two types of experimental studies, based on biochemical isolation methods and raft-perturbation methods. Each entry in the database is annotated by representing its experimental-evidence level with star-symbol (★). The database contains four different experimental evidence levels (labelled level 0, level 1, level 2, and lever 3), that correspond to potentially rafthophilic proteins that were:

(☆☆☆) reported as present in rafts by studies using just a single type of method, without additional validation (level 0).

(★☆☆) identified as present in rafts by two or more biochemical isolation methods (level 1)

(★★☆) down-regulated in two or more raft-perturbation methods (level 2),

(★★★) identified as present in rafts by both the above types of method (level 3),

The positive dataset for training consisted of only human proteins, in total 1118 proteins, that have evidence level of one and three stars; the database currently does not contain any two-star entries for human proteome. The data from other mammals were used for testing the predictive power of the predictor. The negative dataset was retrieved from UniProtKB (UniProt Knowledgebase) (UniProt Consortium, 2018). UniProtKB is a large resource of protein sequences and their detailed annotation. For the negative dataset, we used the reviewed human proteins of UniProt release 2019_08, and we removed the proteins that were already in the positive dataset; this resulted in 19316 proteins.

### Protein features, their retrieval and processing

We labelled protein features as: *general, post-translational modifications* (PTMs), *structural*, and *sequence-motif* and *transmembrane* related. Except for the structural features, the other features are listed in Table 1, 2 and 3, where the source or the way of retrieval are given, too. As specified in detail below, the data for each of the considered features were retrieved either directly from databases, or by available packages and self-written codes.

**Table 2.** **Post-translational modifications as r**etrieved from/by: ^**1**^ UniProt DB, ^**2**^ GPSLipid, ^**3**^ CSS-Palm, and ^**4**^ PhosphoELM DB.

| Post-Translational Modifications | Definition |
| --- | --- |
| Glycosylation<sup>1</sup> | The covalent attachment of a glycan group to a residue. |
| Lipidation<sup>1</sup> | General description of the covalent attachment of lipid groups to a residue. |
| GPI-anchor<sup>1</sup> | The covalent attachment of a GPI-anchor to the C-terminal of a protein. |
| Myristoylation<sup>2</sup> | The covalent attachment of a saturated myristoyl group to the N-terminal glycine residue. |
| Farnesylation<sup>2</sup> | The covalent attachment of a farnesyl group to the C-terminal cysteine residue. |
| Geranyl-geranylation<sup>2</sup> | The covalent attachment of a geranylgeranyl group to the C-terminal cysteine residue. |
| Palmitoylation<sup>3</sup> | The covalent attachment of a saturated palmitic acid to cysteine residue. |
| Phosphorylation<sup>4</sup> | The covalent attachment of a phosphoryl group to a residue. |

**Table 3.** **Transmembrane-protein features**, as retrieved from/by: ^**1**^ UniProt DB, ^**2**^ TMHMM, ^**3**^ Self-written code.

| TM Related Features | Definition |
| --- | --- |
| TMD Length<sup>1</sup> | The number of amino acids in the transmembrane domain of the protein. |
| Accessible Surface Area (ASA)<sup>2</sup> | The surface area of the protein that is accessible to a solvent. |
| Distance between PTMs and TMD<sup>2</sup> | The distance from a PTM location to the TM domain location in a protein. |
| TM type (Single-pass or Multi-pass)<sup>2</sup> | The indication of the type of TM protein |

#### General

With the term general protein-features we refer to those listed in Table 1: sequence length, molecular mass, aliphatic index, net charge, Boman index, isoelectric point, amino acid composition, hydrophobicity, and number of amino acids. Both sequence length and molecular mass (in Da) were retrieved from the UniProtKB database. The length distribution of both positive and negative datasets is shown in Figure 3. We applied a lower limit of 55 residues in order to avoid retrieving data that refers to protein subdomains rather than whole proteins; the upper limit of 5000 residues is applied to remove from the datasets too long proteins and probably multi-protein complexes. Due to these length limitations, 5 and 135 proteins were removed from positive and negative dataset, respectively. Other general features were retrieved using python packages modlAMP (Müller et al., 2017) and BioPython (Cock et al., 2009) or retrieved from UniProtKB. Aliphatic-index, net charge and Boman-index are calculated by a self-written python module employing modlAMP functions (Müller et al., 2017). The aliphatic index (AI) was calculated according to the following formula (Ikai, 1980): *AI* = *X*_*A*_ + *aX*_*V*_ + *b*(*X*_*I*_ + *X*_*L*_), where X_A_, X_V_, X_I_, and X_L_ are mole percent of alanine, valine, isoleucine and leucine, respectively. The coefficients *a* and *b* were calculated as the relative volumes of aliphatic side chains to that of alanine side chains (Ikai, 1980). In addition, the net charge was calculated as the sum of pKa values of amino acids at physiological pH of 7.4, which are determined by the alpha amino group, the alpha carboxyl group and the side chain. The pKa values were obtained from the CRC Handbook of Chemistry and Physics (Haynes, 2015). The Boman-index, as Boman (2003) described it, “is calculated as the sum of the free energies of the respective side chains for transfer from cyclohexane to water and divided by the total number of residues”. Boman-index and aliphatic index are used as density values to train the algorithm.The isoelectric point, amino acid composition, hydrophobicity, number of positive and negative amino acids, and fractions of alpha-helix, beta-sheet and turn structures were calculated by a self-written python module employing Biopython functions. The isoelectric point was calculated in a similar way to the net charge, by using pKa values from the CRC Handbook of Chemistry and Physics (Haynes, 2015). Hydrophobicity was calculated as the sum of hydropathy values of all the amino acids as suggested by Kyte and Doolittle (1982). Data regarding hydrophobicity and amino acid composition features are outputted as density values by the function used.

**Figure 3.**
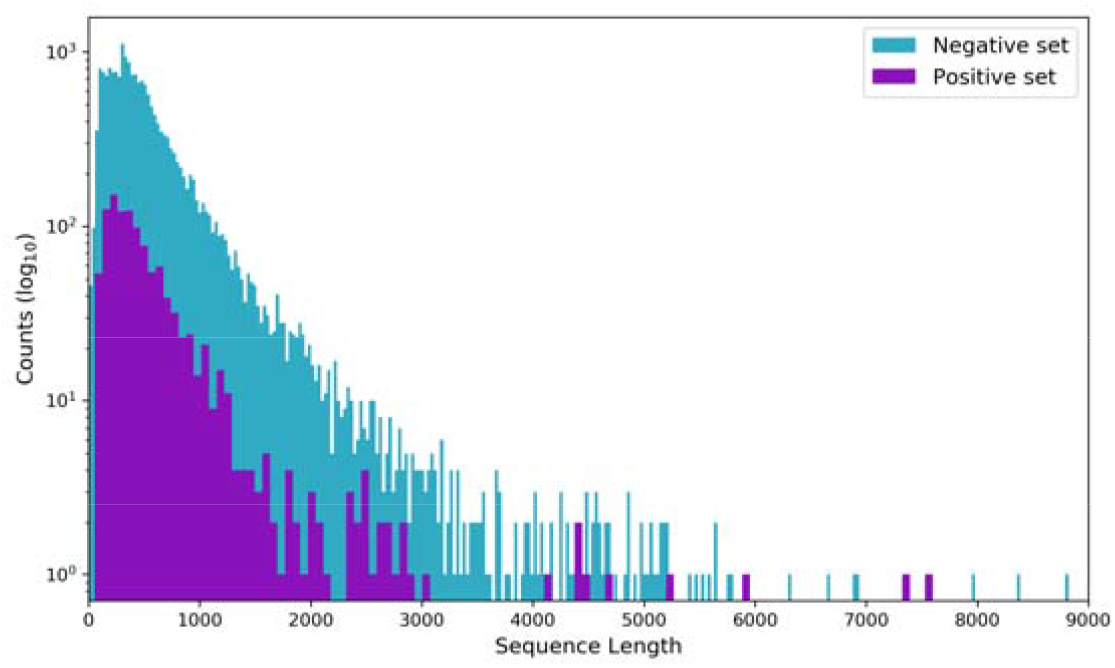
Protein length distribution of the positive and negative datasets. The length of the proteins in the positive dataset ranges from 55 to 7570 amino acids, and for the proteins in the negative dataset has a similar range. Two very long proteins, i.e., 34350 and 14507 amino acid long, were removed from the positive and negative dataset, respectively.

#### Structural

Features regarding *structural* configurations of the proteins include fractions of alpha-helix, beta-sheet and turn structures. The related data were retrieved using BioPython package (Cock et al., 2009) - ProtParam tool or from UniProtKB directly. Secondary structures were calculated from the fractions of the amino acids which tend to be in helix (valine, isoleucine, tyrosine, phenylalanine, tryptophan, and leucine), turn (asparagine, proline, glycine, and serine) or beta-sheet (glutamic acid, methionine, alanine, and leucine). Structural features are presented as density values to the algorithm.

#### Post-translational modifications (PTMs)

PTMs were calculated based on protein sequence information. The list of features is given in Table 2. For myristoylation, farnesylation, geranylgeranylation and palmitoylation we used the local version of online tools: GPS-Lipid (Group-based Prediction System) (Xie et al., 2016) and CSS-Palm (Clustering and Scoring Strategy algorithm) (Ren et al., 2008). We selected threshold ‘high’ for predictions in order to have stringent results. Only the prediction results with a consensus motif were included as possible PTMs and used either as boolean values or counts. Glycosylation, lipidation, and GPI-anchor features are retrieved from UniProt while phosphorylation information is retrieved from PhosphoELM database: a database of experimentally verified phosphorylation sites in eukaryotic proteins (Dinkel et al., 2010).

#### Sequence-motif

We considered CRAC and CARC cholesterol recognition/interaction amino acid sequence motifs, which are listed in Table 1. The CRAC ([L/V]-[X_1-5_]-Y-[X_1-5_]-[K/R]) and CARC ([K/R]-[X_1-5_]-[Y/F]-[X_1-5_]-[L/V]) cholesterol recognition/interaction sequence consensus motifs are expressed in regular expression format. Therefore, we used Python’s regular expression operations (re) module to search the sequences for the motifs, and indicated the existence of the motifs per sequence as either boolean values or counts.

#### Transmembrane

Transmembrane-related features are features specific to protein transmembrane (TM) domains. The list of these features is given in Table 3. TM proteins were retrieved using UniProt annotations and predictions by TMHMM tool (Möller et al., 2001): a method for prediction of transmembrane helices based on a Hidden Markov Model. Combining both results, we identified 439 and 5366 TM proteins in positive and negative sets, respectively. TM domain (TMD) length is retrieved using the TM begin and end locations from UniProtKB annotations and prediction results from TMHMM tool. Similarly, distance from PTMs to TM domain is calculated as the minimum distance from the PTM to the beginning or end of the TM domain. ASA of TM domains is calculated by using the accessible surface areas for amino acid residues when they reside in the membrane (Yuan et al., 2006). Lastly, TM type (single-pass or multi-pass) is also used as an additional feature.

### The XGBoost algorithm

Because of the limited size of the available positive dataset, we used the machine learning algorithm called XGBoost, i.e., the eXtreme Gradient Boosting (Chen and Guestrin, 2016). XGBoost is an algorithm that implements a gradient boosting decision tree framework. Gradient boosting is an ensemble technique where new layers are created to minimize the errors of prior layers using a gradient descent algorithm, and then used altogether to make the final prediction. Decision tree based algorithms, such as XGBoost, are currently considered among the best algorithms for small-to-medium sized datasets.

Prior the implementation of XGBoost, we prepared the datasets by indicating the absence of PTMs and amino acid sequence motifs (CRAC and CARC) with zero, as indicated in Figure 4. Figure 4 shows the percentage of proteins that has at least one PTM or sequence motifs in positive and negative datasets. We then labelled the positive and negative dataset entries as 1 (representing raft proteins) and 0 (representing non-raft proteins), respectively, and down-sampled our negative dataset to have an approximately equal representation of positive and negative samples. Following down-sampling, we merged the data from the positive and the negative dataset, and created the complete dataset that will be used by the XGBoost algorithm. Lastly, the complete dataset is split into training and testing dataset by using %80 and %20 of all data, respectively.

**Figure 4.**
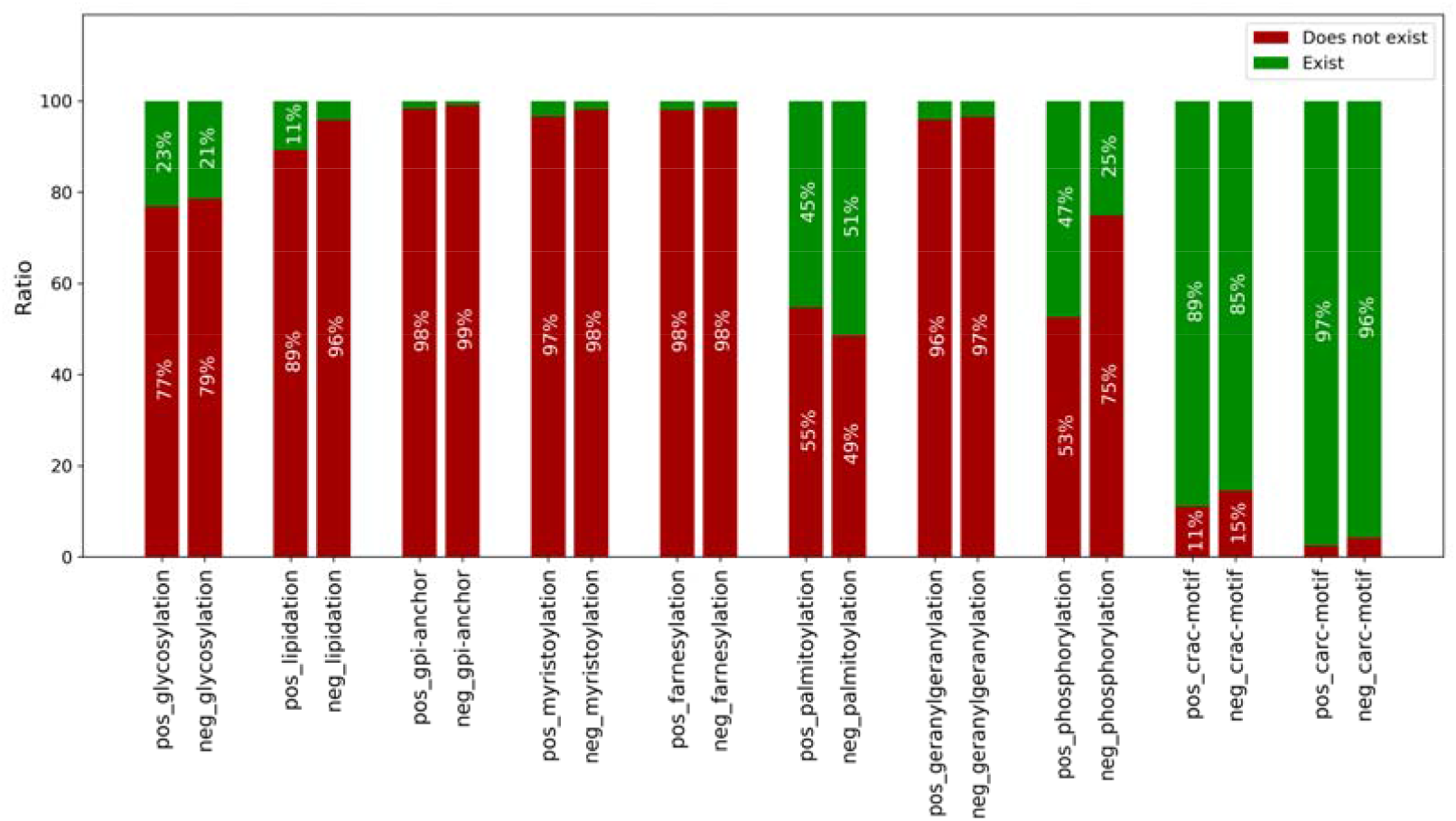
PTM and amino acid sequence motifs abundance in positive and negative datasets. The green bars indicate the percentage of the proteins with one or more PTM, or CRAC/CARC motifs, and the red bars indicate the percentage without any of these features. The first and second bars represent the positive and negative dataset, respectively. In the positive dataset, palmitoylation, phosphorylation, and CRAC and CARC motifs exist in most of the proteins, whereas glycosylation, lipidation, GPI-anchor, myristoylation, farnesylation, and geranylgeranylation hardly occur. This general pattern holds true for the negative dataset as well. A comparison between the two datasets shows that phosphorylation is more abundant in the positive dataset (p<0.001, two sample t-test after Bonferroni correction); this suggests that it could be an important feature for the predictor. Similarly, we found that lipidation, CARC/CRAC motifs, and palmitoylation are also present significantly differently between positive and negative data sets, albeit the difference is less pronounced (p<0.02, two sample t-test after Bonferroni correction).

After preparing the datasets for the XGBoost-based ML algorithm, we first investigated the performance of a baseline and a simplified training on our data. Baseline training was implemented using the ‘DummyClassifier’ class with ‘uniform’ strategy from Sklearn (Pedregosa et al., 2011), which generates random uniform predictions. Simplified classifier was trained as a simplified XGBoost algorithm with default hyperparameters (i.e., the parameters describing the architecture of the predictor). The performance of both methods was then tested on the independent test set, allowing us to create a baseline score, i.e., the performance of the random predictor, and observe the default XGBoost algorithm performance.

### Evaluation Metric

For evaluating the performance of the predictor, we used Area Under the ROC Curve (AUC). The AUC represents the degree of separability - it shows how much the predictor is able to distinguish between classes. The higher the AUC, the better the predictor is at separating different classes. A random predictor would have an AUC of 0.5, whereas a perfect predictor would have an AUC of 1.

## RESULTS AND DISCUSSION

### Effect of feature representation

We first explored the effect of different feature representations on the performance of the predictor. Therefore three different representations for PTMs and sequence motifs features were tested: boolean values, counts, and densities. For this reason, three different datasets were created that represent the features in boolean values (1: if feature exists, 0: if does not), direct counts of PTMs and motifs, and density (counts divided by sequence length). To compare the performance between different feature representations, we trained the machine learning method on the aforementioned datasets and compared their predictive performance on independent test datasets using AUC score as the evaluation metric.

The performance for the three different feature representations are shown in Figure 5. The predictor trained and tested on the density features performs better than the others (Figure 5); this is probably due to the fact that using this representation the information provided to the predictor is not biased by the protein length (e.g., longer proteins would by default have more PTMs).

**Figure 5.**
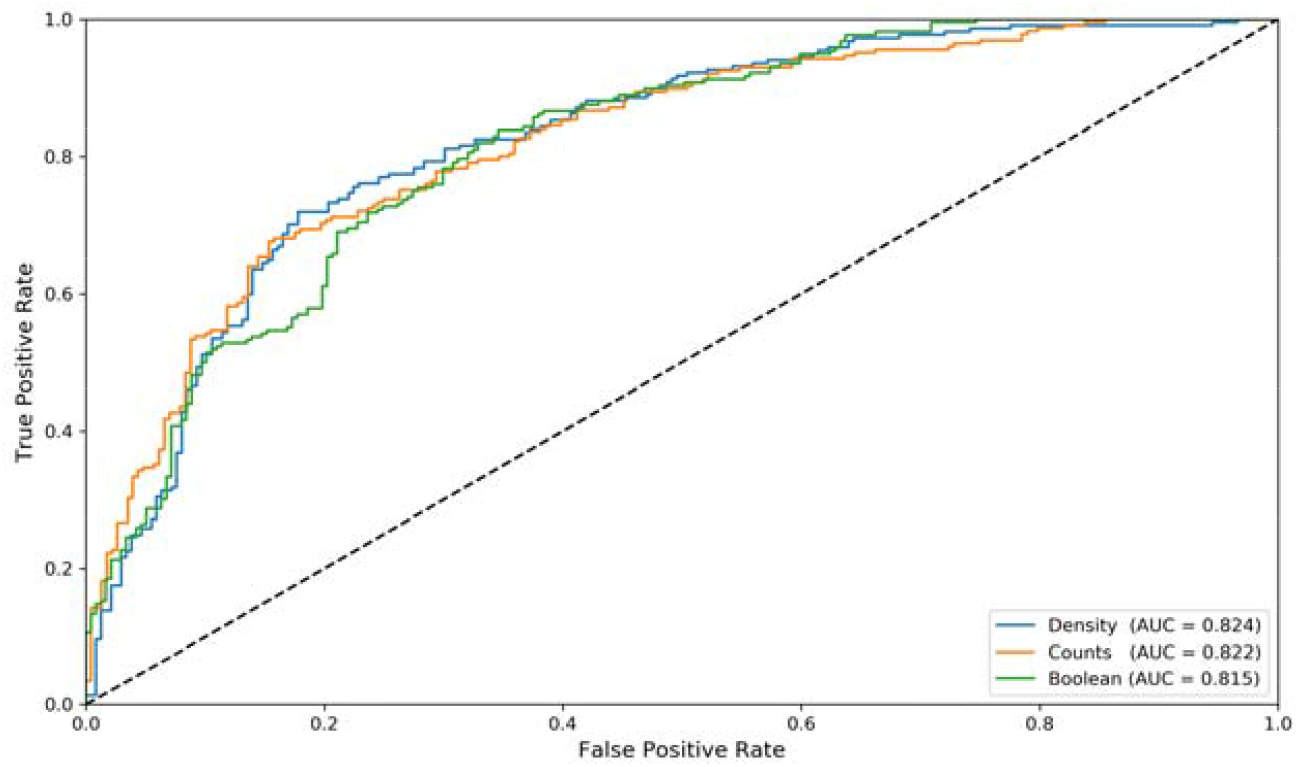
Predictor performance for different feature representations. Performance of three different predictors that are trained and tested with different datasets representing PTM and CRAC/CARC motif features, as boolean, counts, and density (see methods), respectively. Each predictor is trained 100 times, and the median AUC values are used for the comparison. A comparison between the three performance shows that the predictor trained and tested on density values is the best performing one.

### Development of the predictor

After preparing the datasets for the XGBoost-based ML algorithm, we first performed a simplified training with default hyperparameters, and tested its performance on the independent test set. This training allowed us to create a baseline score, i.e., the default performance to be expected from this algorithm. Subsequently, we investigated a broader range of parameters by hyperparameter tuning to further increase the performance of the final predictor. Hyperparameter tuning is the process of searching for the optimal combination of parameters to avoid overfitting. As XGBoost is a decision tree based method, the number of estimators (trees) that are used have an impact on the performance of the predictor. Hence, it is important to use the optimal number of trees to prevent under- or over-fitting. To observe the effect of the number of estimators used, we trained several predictors by changing the number of trees from 1 to 200, and recorded AUC scores for different tree numbers (Supplementary Figure 1). Moreover, we performed hyperparameter tuning for the following possible number of estimators (trees): [35, 45, 50, 70, 80, 90,100].

We used Sklearn’s GridSearchCV (Pedregosa et al., 2011) for hyperparameter tuning, as this approach searches over specified parameter values in an exhaustive way. GridSearchCV uses cross-validation to optimize the results covering two steps of the ML workflow, as shown in Figure 2. To begin with the hyperparameter tuning, we first chose the parameters to be tuned, and some potential values for each of them. Next, we pass these parameters to the GridSearchCV and proceed with the tuning (see Supplementary Table for details on parameters used).

After finalizing the hyperparameter tuning, the tuned predictor was tested on the independent test set. Due to the intrinsic stochasticity of the method and the small size of the dataset, prediction results show some variance. To understand better this variance, we trained the predictor 100 times, each time creating a new train-test split from the complete-set. The overall performance of the predictor, in terms of AUC, is ranging from 0.80 to 0.86 (see Supplementary Figure 2). Considering the stochastic nature of the method, this is an acceptable level of variance as a trade-off to prevent overfitting.

### Protein-feature importance

For training the predictor we have included many features that were either available from the literature or that we calculated. Some features, such as sequence length, molecular mass, amino acid composition, and the positive/negative amino acid counts (see Table 1), are directly derived from the protein sequence, while others, such as aliphatic index, net charge at pH 7.4, isoelectric point, hydrophobicity, and the Boman index are calculated ones (see Materials and Methods for the details).

To identify which features contributed most to the predictions, a drop-column feature importance approach was used (Parr et al., 2018). To this end, we first used all the features for the training to obtain a reference performance score. Next, a feature was dropped, repeated the training with all but one features and a new performance score was calculated. The importance of the feature was then estimated using the difference between the reference performance score and the new performance score. This process was repeated for all the features, resulting in each feature’s overall contribution to the predictor’s performance. The results of the feature importance test over 100 runs (Figure 6) indicate that two of the features have a higher feature importance compared to other features: cysteine fraction and phosphorylation. Cysteine is the residue for palmitoylation modification (Linder, 2001), and it is considered hydrophobic due to its ability to stabilize hydrophobic interactions (Heitmann, 1968). It is likely that the importance of palmitoylation is mainly reflected in the cysteine fraction.

**Figure 6.**
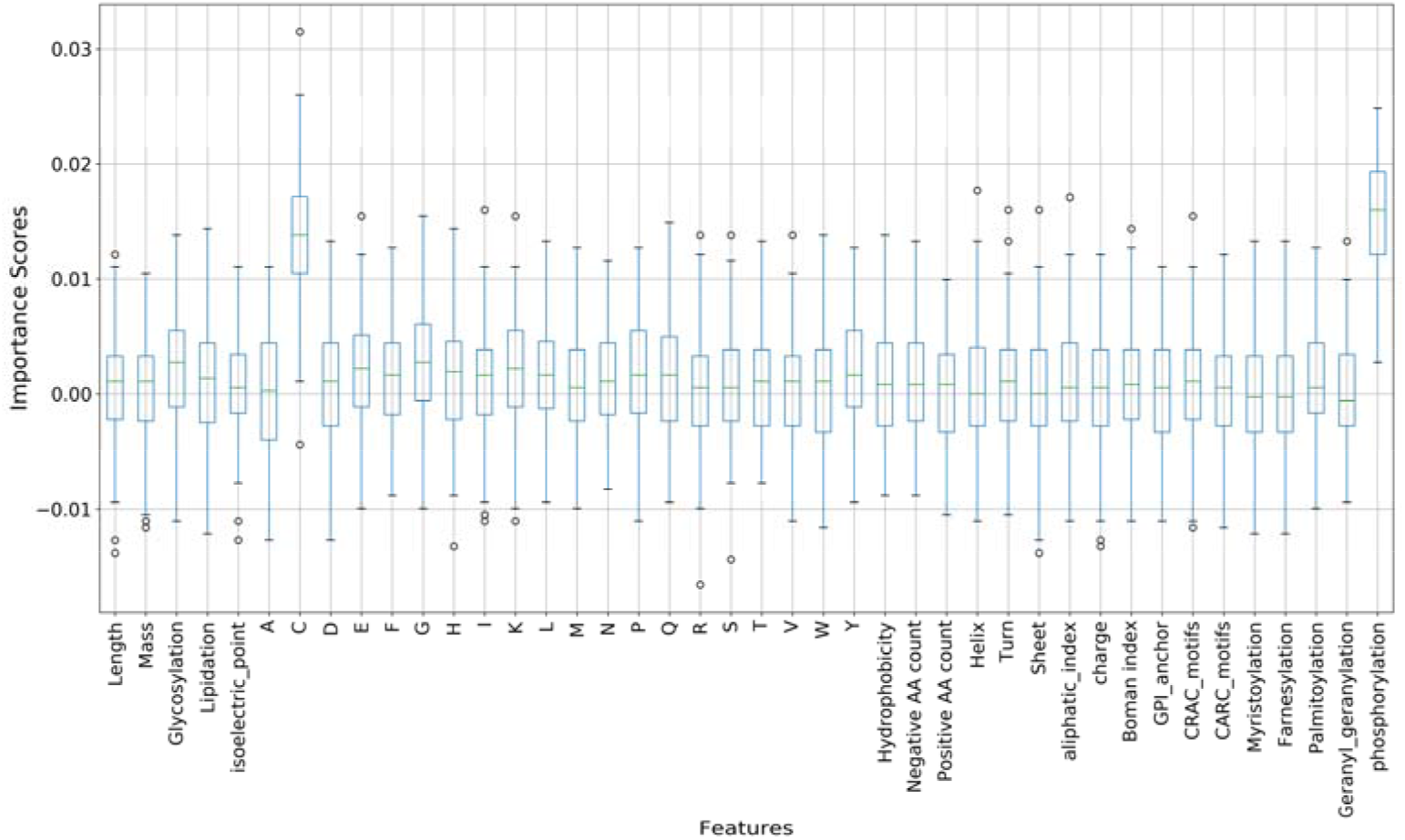
Results of drop-column feature-importance process. Contribution of each feature to the predictor as obtained by training 100 times. Zero line in the plot represents neutral contribution, namely features whose removal/addition would not affect the overall performance. The features with importance above zero are major contributors to the predictor, as removing these would worsen the performance; the features with importance below zero are not contributing to the predictor, and their removal might improve the overall performance. Our results indicated that ProRafts using all the features, and that Cysteine fraction (‘C’) and Phosphorylation are the major contributors.

In previous studies, some of the PTMs were reported to increase the raftophilicity of proteins by mediating lipid-lipid interactions (Lorent and Levental, 2015; Lorent et al., 2017). In our study we also included glycosylation, lipidation, and phosphorylation, to explore their possible importance for protein raftophilicity. Figure 4 (mentioned in Methods) already shows the PTM and amino acid sequence motif abundance in the positive and negative datasets: palmitoylation, and CRAC and CARC motifs occur in most of the proteins (in the positive as well as negative dataset), while glycosylation, lipidation, GPI-anchor, myristoylation, farnesylation, and geranylgeranylation is rare. However, the phosphorylation is more abundant in the positive dataset than the negative one (the amount of phosphorylated proteins in the positive dataset is higher than the negative dataset by %22, *p* ≤ 0.01 to sample t-test), suggesting that it could be relevant in terms of rafthophilicity predictions. This is also clearly visible in Figure 6: although the ProRafts predictor uses most of PTMs features (with the exception of myristoylation, farnesylation, and geranylgeranylation) and cholesterol recognition motifs (CRAC and CARC), phosphorylation is probably the most important feature for raftophilicity. These results are also in accordance with findings from previous investigations, which classify myristoylation as a weak mediator for raft partitioning, and farnesylation and geranylgeranylation as mediators for non-raft membrane domains (Lorent and Levental, 2015; Melkonian et al., 1999).

Multiple signaling molecules are found to associate with lipid rafts, which suggest their role in coordinating signal transduction (Simons and Toomre, 2000). These molecules are activated through phosphorylation and initiate signaling cascades on the cytosolic site of the membrane. This might explain the high contribution of phosphorylation in detecting raft associated proteins. Secondary structure motifs are known to exhibit proteins’ preferences to be in a certain environment, however, only the fraction of turns (median importance score > 0) is used by the predictor to predict raftophilicity. This might be because secondary structure motifs are calculated from physico-chemical properties of amino acids and therefore some of the information about the secondary structure is already embedded in amino acid frequencies.

### Predictor training with different sub-datasets

The positive dataset used above contains both TM and non-TM proteins. As TM proteins exhibit different characteristics compared to the other proteins in the dataset, we studied whether using different datasets for training would improve the performance. To test this, we created three different sub-datasets in addition to the main dataset used above: a sub-dataset with only TM proteins, where specific TMD features (TMD length, ASA of TM domains, distance between PTMs and TMD, and TM type) were used (see Methods), a sub-dataset with only TM proteins but without use of specific TMD features, and a sub-dataset with only non-TM proteins. We trained different predictors using all four datasets, and tested their performance using independent test datasets. In order to compare the results, we plotted the median AUC scores of 100 runs. The results are shown in Figure 7. Interestingly, all four predictors have similarly high performance. Thus, it is not possible to decide which predictor would be the best performing one by looking at these results alone. Transmembrane proteins are embedded in the membrane, hence including TM specific features could improve our predictions. Surprisingly, the performance of TM dataset with and without use of TMD features is very similar. Therefore, addition of the TMD specific features, which contribute to raftophilicity of TM proteins (Lorent et al., 2017), seems not to affect the performance. Unexpectedly, despite having used extra features to predict the raftophilicity of TM proteins, compare to the number of features used for non TM proteins, the performance of the predictor for TM proteins is not higher than the one for the non TM ones. For the raftophilicity predictions of TM proteins we used more features than for the predictions of the non TM ones; unexpectedly, the prediction performance for TM proteins is not higher than the one for the non TM onesFor every model shown in Figure 7, we repeated the feature importance test to identify those features that contributed most to the prediction performance (Supplementary Figure 3 A-C). Due to the different types of proteins being represented in each sub-dataset, the feature importance varies between the predictors. The predictor trained on non-TM raftophilic proteins uses the majority of the features, and Alanine fraction and phosphorylation are the major contributors (Supplementary Figure 3A). On the other hand, the Serine fraction is the least important feature; this might be because Serine residues are used as phosphorylation sites, and therefore the frequency of Serine and the phosphorylation occurrence are not independent. In addition, except from palmitoylation and phosphorylation, no other PTMs contribute to the raftophilicity prediction as was the case for the whole dataset feature importance (compare Figure 6 with Supplementary Figure 3A). In contrast, the predictors trained with TM sub-datasets shows different results regarding feature importance, except that phosphorylation appear to have a major contributor for both. The predictor trained with TM only sub-dataset shows TMD total length and ASA as important features, whereas no other TMD specific feature contributes to the prediction (see Supplementary Figure 3B). The predictor trained with TM only sub-dataset without specific TMD features, on the other hand, has all the features as contributors to differing degrees (see Supplementary Figure 3C).

**Figure 7.**
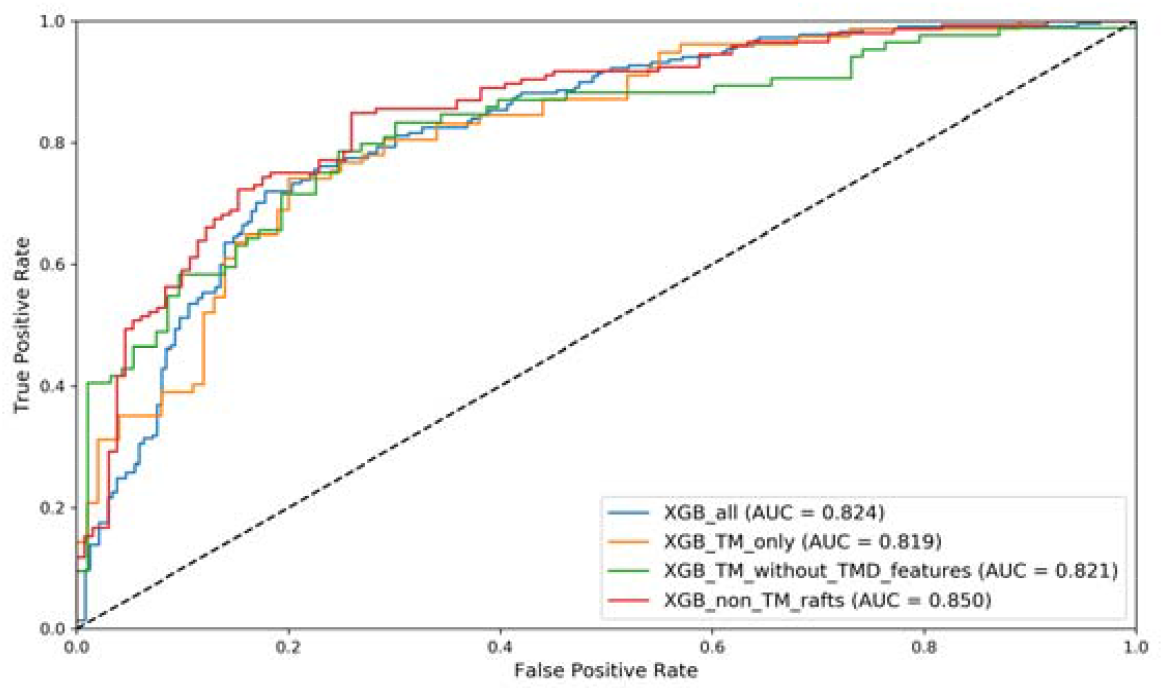
Performance comparison of four different predictors. Four different predictors were trained and tested with their specific datasets. Each predictor is trained 100 times, and the median AUC values are used for the comparison. All four predictors perform similarly high.

### Predictions by ProRafts for other mammal proteins

Finally, we assessed the ProRafts predictive power on other mammal raftophilic proteins, whose data could be retrieved from the RaftPro database. This database classifies raft-proteins according 4 experimental evidence levels, as briefly listed under Methods section. In particular, we retrieved the following datasets from RaftProt database: mouse level 1, 2 and 3; rat level 1 and 3; bovine level 0; mouse level 0; and rat level 0. For these raftophilic proteins we “calculated” the features and tested the XGBoost predictor trained on the human level 1 and 3 entries.

Figure 8 shows that the performance (more than %80) on mouse proteins (level 1, 2, and 3) and rat proteins (level 1 and 3), which is remarkably similar to the performance on human proteins level 1 and 3 (see Fig. 7, XGB_all, blue line, and Supplementary Figure 2). In addition, %57 of the proteins in the rat level 0 dataset are predicted to be raftophilic, whereas mouse level 0 prediction results are a bit lower (%47). Furthermore, %70 of the bovine level 0 entries are predicted to be raftophilic, which is surprising because the prediction rate for other level 0 mammals is much lower. We cannot exclude the possibility that the small amount of available validated raftophilic proteins in the training dataset could make the predictor insensitive to certain features that have predictive power, but are not sampled adequately in the training dataset. Nevertheless, ProRafts trained on only human entries is able to classify other mammal (mouse, rat and bovine) raftophilic proteins. This suggests that raftophilicity prediction is not human specific, and our model can also be used for predicting other mammal raftophilic proteins.

**Figure 8.**
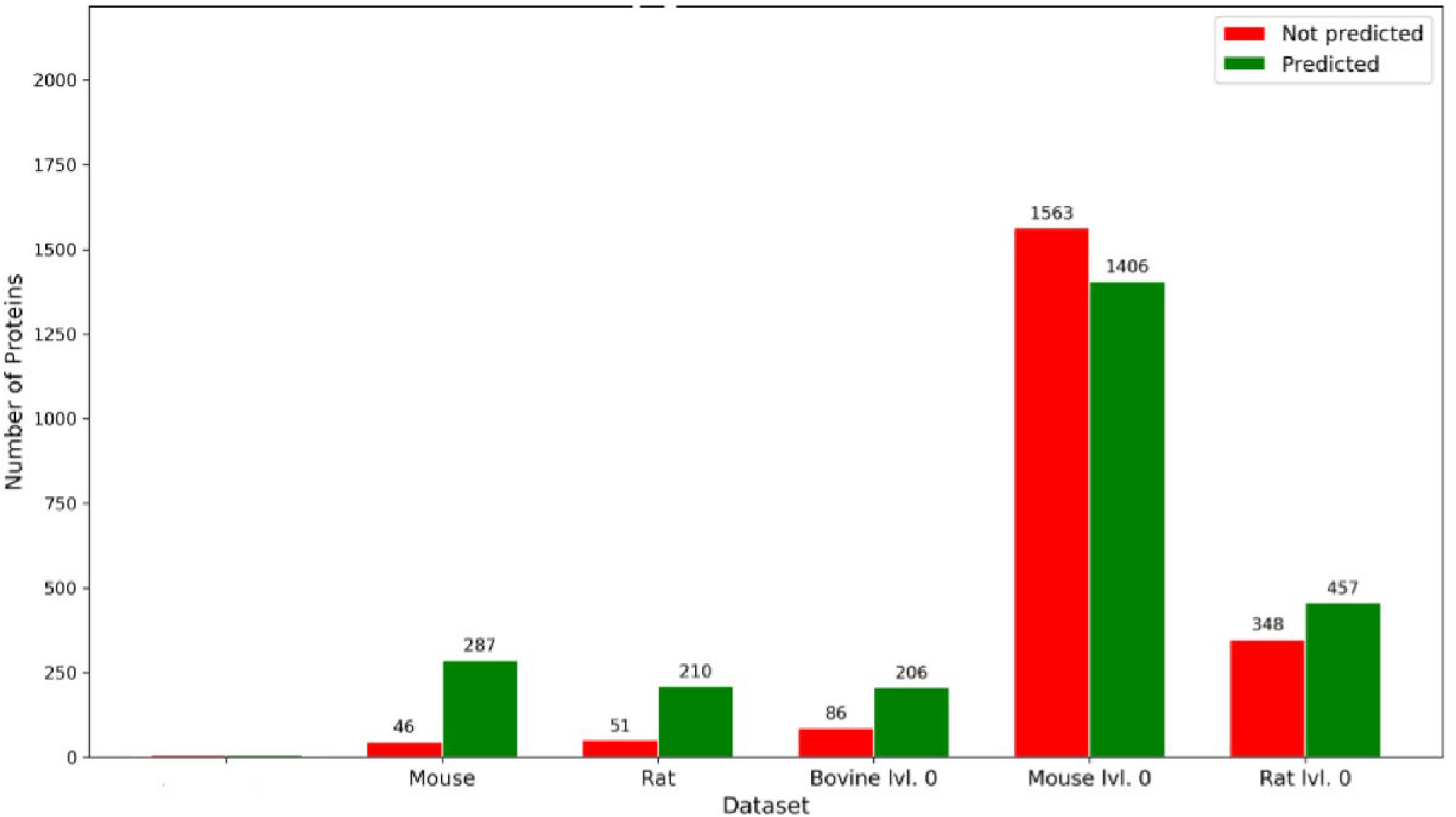
ProRafts predictive performance for other mammal raft proteins. The performance of the best predictor from Figure 7 (XGB_all, blue line), which is trained on human level 1 and 3 entries from RaftProt database, is tested on five different datasets. ProRafts is able to predict successfully raft proteins from other mammals (rats, mouse and bovine).

## CONCLUSION

The term *raftophilicity* was used (Simons and Sampaio, 2011) to indicate the affinity of proteins for biomembranes’ lipid domains, called ‘rafts’. We developed a Machine Learning method, and a resulting predictor, ProRafts, via which it is possible to predict mammal-protein raftophilicity with a reasonable high accuracy—despite the fact that the data that we could retrieve from available databases resulted in a limited size training dataset of human proteins denoted as raftophilic. Interestingly, ProRafts performed well with regards to other mammals’ proteins, e.g., mouse- and rat-proteins, suggesting that it can identify relevant features of raft proteins in mammals. Therefore, ProRafts can be used in raftophilicity studies of other mammals than humans. In addition, our results indicate that, raftophilicity-wise, some proteins features are more significant than others; this is especially the case for phosphorylation, which may play a more relevant role for protein raftophilicity than indicated by previous studies.

ProRafts predictions’ performance depends on the validity and size of the training dataset used. The predictions presented in this paper are based on the assumption that proteins in the dataset used for training the predictor are indeed raftophilic, as stated in the database from raft-proteins, RaftProt V2, from where the data that we used were retrieved. Data from validated proteins that will become available in the future will increase the size of the training dataset, and hence will improve the ProRafts predictor performance. Nevertheless, the predictions by actual ProRafts can be used as a guideline for the identification of human proteins with unknown raftophilicity: once the proteins are identified as potentially raftophilic by ProRafts, and their raftophilicity is sub-sequentially validated by experimental methods, these proteins could then be used as markers for diseases, or could serve as targeting sites for anti-disease therapeutics. Based on evidences that rafts play a role in the interaction between viruses and host cells—herewith in binding, endocytosis, and replication of viruses, including the corona viruses—a recent datamining study (Yurtsever and Lorent, 2020), compared structural properties of proteins from rafts and viruses, and provided some insight that could be useful for targeting antiviral compounds. A tool such as ProRafts can facilitate similar raft-related datamining investigations.

We tackled the protein raftophilicity prediction task as a categorical classification one; the outcome is one of the two output categories, “raftophilic” or “non-raftophilic”, and the identification of those protein features that may contribute significantly to raftophilicity. However, protein-raftophilicity should be viewed as a propensity for the proteins to partition between raft domains and non-raft domains, which are not static entities. Therefore protein partitioning into rafts depends on the dynamic changes to which biomembranes are subjected during the life cycle of cells. To account for this dependence one should estimate the likelihood of partitioning (Silvius, 2005), and then use a regression method—rather than a categorical classification method, as we did—where the raftophilicity of a given protein would be outputted as a numerical value, which can then be converted to a ratio of partitioning. However, this would require quantitative data on raft partitioning, and to our knowledge these data are still sparse.

Our next step, in the development of a new version of ProRafts, will be to train the predictor with additional features, such as the sphingolipid-binding sequence homologous motifs, which were identified in the human immunodeficiency virus (HIV)-1 surface envelope glycoprotein gp120, the human prior proteins (PrP) and the Alzheimer β-amyloid peptide (Mahfoud et al., 2002; Fantini, 2003), and other mammalian membrane proteins (Björkholm et al., 2014).

## Availability

The ProRafts predictor is downloadable as source code (in Python) from the github repository https://github.com/DenizY7/ProRafts. The datasets for training/testing the predictor are also available from the same depository.

## Authors’s Contributions

DY developed the predictor under guidance of CK. MMS initiated the project of building a raftophilicity predictor based on protein features, reviewed biophysical/biological aspects of the project, and wrote a substantial part of the manuscript. Both DY and CK contributed to the manuscript preparation.

## Acknowledgements

We thank QIMR Berghofer Medical Research Institute for the RaftProt database and UniProt Consortium for the UniProt Knowledgebase, from which we retrieved the datasets for this study. We thank EMBL Heidelberg for the PhosphoELM database, from which we retrieved the information regarding phosphorylation. In addition, we thank the Technical University of Denmark (DTU) for the TMHMM tool, and the CUCKOO Workgroup (China) for the GPSLipid and CSS-Palm tools. CK and DY, thanks Joseph H. Lorent, Utrecht University (The Netherland) for enlightening discussions. The idea of using a bioinformatics approach to predict protein raftophilicity originated from fruitful discussions that one of us, MMS, had with scientists at the former Center for Biological Sequence Analysis at DTU; the present work was inspired by the outcome of those discussions.

## REFERENCES

Antonini A, Caioli S, Saba L, Vindigni G, Biocca S, Canu N, Zona C. Membrane cholesterol depletion in cortical neurons highlights altered NMDA receptor functionality in a mouse model of amyotrophic lateral sclerosis. Biochim Biophys Acta Mol Basis Dis. 1864, 509–519 (2018). Epub 2017 Nov 24. https://doi.org/10.1016/j.bbadis.2017.11.008

Baglivo M, Baronio M, Natalini G, Beccari T, Chiurazzi P, Fulcheri E, Petralia P, Michelini S, Fiorentini G, Miggiano G, Morresi A, Tonini G, and Bertelli M. Natural small molecules as inhibitors of coronavirus lipid-dependent attachment to host cells: a possible strategy for reducing SARS-COV-2 infectivity? Acta Bio-medica: Atenei Parmensis 91, 161–164 (2020). https://doi.org/10.23750/abm.v91i1.9402

Baier CJ, Fantini J, Barrantes FJ. Disclosure of cholesterol recognition motifs in transmembrane domains of the human nicotinic acetylcholine receptor. Sci Rep. 1, 69–76 (2011). https://doi.org/10.1038/srep00069

Bhatia T, Cornelius F, Ipsen JH. Biochem. Exploring the raft-hypothesis by probing planar bilayer patches of freestanding giant vesicles at nanoscale resolution, with and without Na,K-ATPase. Biochim. Biophys. Acta 1858: 3041–3049 (2016). https://doi.org/10.1016/j.bbamem.2016.09.001

Björkholm, P, Ernst AM, Hacke M, Wieland F, Brügger B, von Heijne G. Identification of novel sphingolipid-binding motifs in mammalian membrane proteins. Biochim. Biophys. Acta 1838, 2066–2070 (2014). https://doi.org/10.1016/j.bbamem.2014.04.026

Boman H. Antibacterial peptides: basic facts and emerging concepts. J. Internal Medicine 254, 197–215 (2003). https://doi.org/10.1046/j.1365-2796.2003.01228.x

Carquin M, D’Auria L, Pollet H, Bongarzone ER, Tyteca D. Recent progress on lipid lateral heterogeneity in plasma membranes: From rafts to submicrometric domains. Progress in Lipid Research 62, 1–24 (2016). https://doi.org/10.1016/j.plipres.2015.12.004

Cartocci V, Servadio M, Trezza V, Pallottini V. Can cholesterol metabolism modulation affect brain function and behavior? J. Cell. Physiol. 232, 281–286 (2017). https://doi.org/10.1002/jcp.25488

Cheng X, Smith JC. Biological membrane organization and cellular signaling. Chemical Reviews 119, 5849–5880 (2019). https://doi.org/10.1021/acs.chemrev.8b00439

Chen T, Guestrin C. XGBoost: A scalable tree boosting system. In Proceedings of the 22nd ACM SIGKDD International Conference on Knowledge Discovery and Data Mining, pp. 785–794, Association for Computing Machinery (2016). https://doi.org/10.1145/2939672.2939785

Cock PJ, Antao T, Chang JT, Chapman BA, Cox CJ, Dalke A, Friedberg I, Halmerick T, Kauff F, Wilczynski B, de Hoon MJL. Biopython: freely available Python tools for computational molecular biology and bioinformatics. Bioinformatics 25, 1422–1423 (2009). https://doi.org/10.1093/bioinformatics/btp163

Colin J, Gregory-Pauron L, Lanhers M-C, Claudepierre T, Corbier C, Yen FT, Malaplate-Armand C, Oster T. Membrane raft domains and remodeling in aging brain. Biochimie, 130, 178–187 (2016). https://doi.org/10.1016/j.biochi.2016.08.014

Díaz ML. Membrane physiology and biophysics in the next decade: an open balcony to multiple scenarios. Frontiers in Physiology 1, 23 (2010). https://doi.org/10.3389/fphys.2010.00023

Díaz L, Fabelo N, Ferrer I, Marín R. “Lipid raft aging” in the human frontal cortex during nonpathological aging: gender influences and potential implications in Alzheimer’s disease. Neurobiol. Aging 67, 42–52 (2018). https://doi.org/10.1016/j.neurobiolaging.2018.02.022

Di Scala C, Baier CJ, Evans LS, Williamson PTF, Fantini J, Barrantes FJ. Relevance of CARC and CRAC Cholesterol-Recognition Motifs in the Nicotinic Acetylcholine Receptor and Other Membrane-Bound Receptors. In Current Topics in Membranes, Ed. I. Levitan, Academic Press, Volume 80, Chapter One, Pages 3-23, ISSN 1063-5823, ISBN 9780128093887 (2017) https://doi.org/10.1016/bs.ctm.2017.05.001

Diaz-Rohrer B, Levental KR, Levental I. Rafting through traffic: Membrane domains in cellular logistics. Biochim Biophys Acta. 1838, 3003–3013 (2014a). Epub 2014 Aug 15. Review. https://doi.org/10.1016/j.bbamem.2014.07.029

Diaz-Rohrer BB, Levental KR, Simons K, Levental I. Membrane raft association is a determinant of plasma membrane localization. Proc Natl Acad Sci U S A 111, 8500–8505 (2014b). Epub 2014 May 27. https://doi.org/10.1073/pnas.1404582111

Dinkel H, Chica C, Via A, Gould CM, Jensen LJ, Gibson TJ, Diella F. Phospho. ELM: a database of phosphorylation sites—update 2011. Nucleic Acids Res. 39(suppl_1), D261–D267 (2010). https://doi.org/10.1093/nar/gkq1104

Douglass AD, Vale RD. Single-molecule microscopy reveals plasma membrane microdomains created by proteinprotein networks that exclude or trap signaling molecules in T cells. Cell 121, 937–950 (2005). https://doi.org/10.1016/j.cell.2005.04.009

Epand RM. Cholesterol and the interaction of proteins with membrane domains. Prog Lipid Res 45, 279–294 (2006). https://doi.org/10.1016/j.plipres.2006.02.001

Epand RM, Thomas A, Brasseur R, Epand RF. Cholesterol interaction with proteins that partition into membrane domains: an overview. In Cholesterol Binding and Cholesterol Transport Proteins. Vol. 51, pp. 253–278. Dordrecht: Springer (2010). https://doi.org/10.1007/978-90-481-8622-8_9

Estep T, Mountcastle D, Barenholz Y, Biltonen R, Thompson T. Thermal behavior of synthetic sphingomyelin-cholesterol dispersions. Biochemistry 18, 2112–2117 (1979). https://doi.org/10.1021/bi00577a042

Fantini J. How sphingolipids bind and shape proteins: molecular basis of lipid-protein interactions in lipid shells, rafts and related biomembrane domains. Cellular and Molecular Life Sciences CMLS 60, 1027–1032 (2003). https://doi.org/10.1007/s00018-003-3003-1

Fantini J, Barrantes FJ. Sphingolipid/cholesterol regulation of neurotransmitter receptor conformation and function. Biochim. Biophys. Acta 1788, 2345-2361 (2009). https://doi.org/10.1016/j.bbamem.2009.08.016

Fantini J, Barrantes FJ. How cholesterol interacts with membrane proteins: an exploration of cholesterol-binding sites including CRAC, CARC, and tilted domains. Frontiers in Physiology 4, 1–9 (2013). https://doi.org/10.3389%2ffphys.2013.00031

Fantini J, Di Scala C, Evans LS, Williamson PT, Barrantes FJ. A mirror code for protein-cholesterol interactions in the two leaflets of biological membranes. Sci. Rep. 6, 21907 (2016). https://doi.org/10.1038/srep21907

Foster LJ, Chan QW. Lipid raft proteomics: more than just detergent-resistant membranes. Subcell Biochem. 43, 35–47 (2007). https://doi.org/10.1007/978-1-4020-5943-8_4

Giese SI, Woerz I, Homann S, Tibroni N, Geyer M, Fackler OT. Specific and distinct determinants mediate membrane binding and lipid raft incorporation of HIV-1(SF2) Nef. Virology. 355, 175–191 (2006). Epub 2006 Aug 17. https://doi.org/10.1016/j.virol.2006.07.003

Goodsaid-Zalduondo F, Rintoul D, Carlson J, Hansel W. Luteolysis-induced changes in phase composition and fluidity of bovine luteal cell membranes. Proc. Nat. Acad. Sci. USA 79, 4332–4336 (1982). https://doi.org/10.1073/pnas.79.14.4332

Guo H, Huang M, Yuan Q, Wei Y, Gao Y, Mao L, Gu L, Tan YW, Zhong Y, Liu D, Sun S. The Important Role of Lipid Raft-Mediated Attachment in the Infection of Cultured Cells by Coronavirus Infectious Bronchitis Virus Beaudette Strain. PLoS ONE 12(1):e0170123 (2017) https://doi.org/10.1371/journal.pone.0170123

Hawkins DM. The problem of overfitting. J. Chem. Inf. Comput. Sci. 44, 1–12 (2004). https://doi.org/10.1021/ci0342472

Haynes WM. CRC Handbook of Chemistry and Physics. 96th edition, CRC press (2015).

Heitmann P. A model for sulfhydryl groups in proteins. Hydrophobic interactions of the cysteine side chain in micelles. Eur. J. Biochemistry 3, 346–350 (1968). https://doi.org/10.1111/j.1432-1033.1968.tb19535.x

Hryniewicz-Jankowska A, Augoff K, Biernatowska A, Podkalicka J, Sikorski AF. Membrane rafts as a novel target in cancer therapy. Biochim. Biophys. Acta 1845, 155–165 (2014). https://doi.org/10.1016/j.bbcan.2014.01.006

Ikai A. Thermostability and aliphatic index of globular proteins. J. Biochem, 88, 1895–1898 (1980). https://doi.org/10.1093/oxfordjournals.jbchem.a133168

Ipsen JH, Karlström G, Mouritisen OG, Wennerström H, Zuckermann MJ. Phase-equilibria in the phosphatidylcholine-cholesterol system. Biochim. Biophys. Acta, 905, 162–172 (1987). https://doi.org/10.1016/0005-2736(87)90020-4

Israelachvili J, Marčelja S, Horn RG. Physical principles of membrane organization. Quart. Rev. Biophys. 13, 121–200 (1980). https://doi.org/10.1017/S0033583500001645

Jacobson K, Mouritsen OG, Anderson RG. Lipid rafts: at a crossroad between cell biology and physics. Nat. Cell Biol. 9, 7–14 (2007). https://doi.org/10.1038/ncb0107-7

Jensen LJ, Gupta R, Blom N, Devos D, Tamames J, Keşmir C, Nielsen H, Staerfeldt HH, Rapacki K, Workman C, Andersen CA, Knudsen S, Krogh A, Valencia A, Brunak S. Prediction of human protein function from post-translational modifications and localization features. J. Mol. Biol. 319, 1257–1265 (2002). https://doi.org/10.1016/S0022-2836(02)00379-0

Karnovsky MJ, Kleinfeld AM, Hoover RL, Klausner RD. The concept of lipid domains in membranes. J. Cell Biol. 94, 1–6 (1982). https://doi.org/10.1083/jcb.94.1.1

Kenworthy AK. Have we become overly reliant on lipid rafts? Talking Point on the involvement of lipid rafts in T-cell activation. EMBO Rep., 9, 531–535 (2008). https://doi.org/10.1038/embor.2008.92

Korade Z, Kenworthy AK. Lipid rafts, cholesterol, and the brain. Neuropharmacology 55, 1265–1273 (2008). https://doi.org/10.1016/j.neuropharm.2008.02.019

Kyte J, Doolittle RF. A simple method for displaying the hydropathic character of a protein. J. Mol. Biol. 157, 105–132 (1982). https://doi.org/10.1016/0022-2836(82)90515-0

Lajoie P, Nabi IR. Regulation of raft-dependent endocytosis. J. Cell. Mol. Med. 11, 644–653 (2007). https://doi.org/10.1111/j.1582-4934.2007.00083.x

Levental I. Lipid rafts come of age. Nat. Rev. Mol. Cell Biol. 21, 420 (2020). https://doi.org/10.1038/s41580-020-0252-y

Levental I, Levental KR, Heberle FA. Lipid Rafts: Controversies Resolved, Mysteries Remain. Trends Cell Biol. 30, 341–353 (2020) https://doi.org/10.1016/j.tcb.2020.01.009

Levental I, Lingwood D, Grzybek M, Coskun U, Simons K. Palmitoylation regulates raft affinity for the majority of integral raft proteins. Proc. Natl. Aca.d Sci. USA 107, 22050–22054 (2010a). https://doi.org/10.1073/pnas.1016184107

Levental I, Grzybek M, Simons K. Greasing their way: lipid modifications determine protein association with membrane rafts. Biochemistry 49, 6305–6316 (2010b). https://doi.org/10.1021/bi100882y

Li H, Papadopoulos V. Peripheral-type benzodiazepine receptor function in cholesterol transport. Identification of a putative cholesterol recognition/interaction amino acid sequence and consensus pattern. Endocrinology 139, 4991–4997 (1998). https://doi.org/10.1210/endo.139.12.6390

Lin Q, E. London. Altering Hydrophobic Sequence Lengths Shows That Hydrophobic Mismatch Controls Affinity for Ordered Lipid Domains (Rafts) in the Multitransmembrane Strand Protein Perfringolysin O. J. Biol. Chem. 288, 1340–1352 (2013). https://doi.org/10.1074/jbc.M112.415596

Linder ME. 8 Reversible modification of proteins with thioester-linked fatty acids. Enzym. 21, 215–240 (2001). https://doi.org/10.1016/S1874-6047(01)80021-4

Listowski MA, Leluk J, Kraszewski S, Sikorski AF. Cholesterol Interaction with the MAGUK Protein Family Member, MPP1, via CRAC and CRAC-Like Motifs: An In Silico Docking Analysis. Deschenes RJ, ed. PLoS ONE 10(7):e0133141 (2015). https://doi.org/10.1371/journal.pone.0133141

Lorent JH, Levental I. Structural determinants of protein partitioning into ordered membrane domains and lipid rafts. Chem. Phys. Lipids, 192, 23–32 (2015). https://doi.org/10.1016/j.chemphyslip.2015.07.022

Lorent JH, Diaz-Rohrer B, Lin X, Spring K, Gorfe AA, Levental KR., and Levental, I. Structural determinants and functional consequences of protein affinity for membrane rafts. Nat. Commun. 8, 1219 (2017). https://doi.org/10.1038/s41467-017-01328-3

Lorizate M, Kräusslich H-G. Role of Lipids in Virus Replication. Cold Spring Harb Perspect Biol 2011;3:a004820 (2011). https://doi.org/10.1101/cshperspect.a004820

Magee AI, Parmryd I. Detergent-resistant membranes and the protein composition of lipid rafts. Genome Biol. 4, 234- (2003). https://doi.org/10.1186/gb-2003-4-11-234

Mahfoud R, Garmy N, Maresca M, Yahi N, Puigserver A, Fantini J. Identification of a common sphingolipid-binding domain in Alzheimer, prion, and HIV-1 proteins. J. Biol. Chem. 277, 11292–11296 (2002). https://doi.org/10.1074/jbc.M111679200

Marin R, Rojo JA, Fabelo N, Fernandez CE, Diaz M. Lipid raft disarrangement as a result of neuropathological progresses: a novel strategy for early diagnosis? Neuroscience, 245, 26–39 (2013). https://doi.org/10.1016/j.neuroscience.2013.04.025

Melkonian KA, Ostermeyer AG, Chen JZ, Roth MG, Brown DA. Role of Lipid Modifications in Targeting Proteins to Detergent-resistant Membrane Rafts Many raft proteins are acylated, while few are prenylated. J. Biol. Chem. 274, 3910–3917 (1999). https://doi.org/10.1074/jbc.274.6.3910

Mohamed A, Shah AD, Chen D, Hill MM. RaftProt V2: understanding membrane microdomain function through lipid raft proteomes. Nucleic Acids Res. 47(D1), D459–D463 (2019). https://doi.org/10.1093/nar/gky948

Munro S. An investigation of the role of transmembrane domains in Golgi protein retention. EMBO J. 14, 4695–4704 (1995). https://doi.org/10.1002/j.1460-2075.1995.tb00151.x

Möller S, Croning MD, Apweiler R. Evaluation of methods for the prediction of membrane spanning regions. Bioinformatics 17, 646–653 (2001). https://doi.org/10.1093/bioinformatics/17.7.646

Müller AT, Gabernet G, Hiss JA., Schneider G. modlAMP: Python for antimicrobial peptides. Bioinformatics, 33, 2753–2755 (2017). https://doi.org/10.1093/bioinformatics/btx285

Nicolson GL. Transmembrane control of the receptors on normal and tumor cells: I. Cytoplasmic influence over cell surface components. Biochim. Biophys. Acta 457, 57–108 (1976). https://doi.org/10.1016/0304-4157(76)90014-9

Nicolson GL. The Fluid-Mosaic Model of Membrane Structure: Still relevant to understanding the structure, function and dynamics of biological membranes after more than 40 years. Biochim. Biophys. Acta 1838, 1451–1466 (2014). https://doi.org/10.1016/j.bbamem.2013.10.019

Owen DM, Williamson DJ, Magenau A, Gaus K. Sub-resolution lipid domains exist in the plasma membrane and regulate protein diffusion and distribution. Nat. Commun. 3,1256 (2012). https://doi.org/10.1038/ncomms2273

Parton RG, Simons K. The multiple faces of caveolae. Nat. Rev. Mol. Cell. Biol. 8, 185–194 (2007). https://doi.org/10.1038/nrm2122

Parr T, Turgutlu K, Csiszar C, Howard J. Beware Default Random Forest Importances. Online article https://explained.ai/rf-importance/index.html (2018).

Pedregosa F, Varoquaux G, Gramfort A, Michel V, Thirion B, Grisel O, Blondel M, Prettenhofer P, Weiss R, Dubourg, V, Vanderplas J, Passos A, Cournapeau D, Brucher M, Perrot M, Duchesnay E. Scikit-learn: Machine learning in Python. J. Machine Learning Res. 12, 2825–2830 (2011). https://arxiv.org/abs/1201.0490

Pike LJ. Lipid rafts: heterogeneity on the high seas. Biochem. J. 378, 281–292 (2004). https://doi.org/10.1042/bj20031672

Pike LJ. Rafts defined: a report on the Keystone Symposium on lipid rafts and cell function. J. Lipid Res. 47, 1597–1598 (2006). https://doi.org/10.1194/jlr.E600002-JLR200

Quinn PJ, Wolf C. The liquid-ordered phase in membranes. Biochim. Biophys. Acta 1788, 33–46 (2009). https://doi.org/10.1016/j.bbamem.2008.08.005

Rabilloud T. Membrane proteins and proteomics: love is possible, but so difficult. Electrophoresis 30 (Suppl. 1), S174–S180 (2009). https://doi.org/10.1002/elps.200900050

Ren J, Wen L, Gao X, Jin C, Xue Y, Yao X. CSS-Palm 2.0: an updated software for palmitoylation sites prediction. Protein Eng. Des. Sel. 21, 639–644 (2008). https://doi.org/10.1093/protein/gzn039

Santos G, Díaz M, Torres NV. Lipid raft size and lipid mobility in non-raft domains increase during aging and are exacerbated in APP/PS1 mice model of Alzheimer’s disease. Predictions from an agent-based mathematical model. Front. Physiol. 7:90 (2016). https://doi.org/10.3389/fphys.2016.00090

Schengrund C-L. Lipid rafts: Keys to neurodegeneration. Brain Research Bulletin 82, 7–17 (2010). https://doi.org/10.1016/j.brainresbull.2010.02.013

Schug ZT, Frezza C, Galbraith LC, Gottlieb E. The music of lipids: How lipid composition orchestrates cellular behaviour. Acta Oncologica 51, 301–310 (2012). https://doi.org/10.3109/0284186X.2011.643823

Semrau T, Schmidt T. Membrane heterogeneity – from lipid domains to curvature effects. Soft Matter 5, 3174–3186 (2009). https://doi.org/10.1039/B901587F

Sengupta P, Seo AY, Pasolli HA, Song YE, Johnson MC, Lippincott-Schwartz J. A lipid-based partitioning mechanism for selective incorporation of proteins into membranes of HIV particles. Nat Cell Biol. 21, 452–461 (2019). Epub 2019 Apr 1. Erratum in: Nat Cell Biol. 2019 Apr 10. https://doi.org/10.1038/s41556-019-0300-y

Sezgin E, Levental I, Mayor S, Eggeling C. The mystery of membrane organization: composition, regulation and roles of lipid rafts. Nat. Rev. Mol. Cell Biol., 18(6), 361–374 (2017). https://doi.org/10.1038/nrm.2017.16

Sharpe HJ, Stevens TJ, Munro S. A comprehensive comparison of transmembrane domains reveals organelle-specific properties. Cell 142, 158–169 (2010). https://doi.org/10.1016/j.cell.2010.05.037

Silvius JR. Partitioning of membrane molecules between raft and non-raft domains: Insights from model-membrane studies. Biochim. Biophys. Acta 1746, 193–202 (2005). https://doi.org/10.1016/j.bbamcr.2005.09.003

Simons K, Ikonen E. Functional rafts in cell membranes. Nature 387, 569–572 (1997) https://doi.org/10.1038/42408

Simons K, Gerl MJ. Revitalizing membrane rafts: new tools and insights. Nat. Rev. Mol. Cell. Biol. 11, 688–699 (2010). https://doi.org/10.1038/nrm2977

Simons K, Sampaio JL. Membrane organization and lipid rafts. Cold Spring Harb Perspect Biol. 2011;3:a004697 (2011). https://doi.org/10.1101/cshperspect.a004697

Simons K, Toomre D. Lipid rafts and signal transduction. Nat. Rev. Mol. Cell Biol., 1, 31–39 (2000). https://doi.org/10.1038/35036052

Singer SJ, Nicolson GL. The Fluid Mosaic Model of the Structure of Cell Membranes. Science 175, 720–731 (1972). https://doi.org/10.1126/science.175.4023.720

Sodt AJ, Sandar ML, Gawrisch K, Pastor RW, Lyman E. The Molecular Structure of the Liquid-Ordered Phase of Lipid Bilayers. J. Am. Chem. Soc. 136, 725–732 (2014). https://doi.org/10.1021/ja4105667

Staubach S, Hanisch F-G. Lipid rafts: signaling and sorting platforms of cells and their roles in cancer. Expert Review of Proteomics 8, 263–277 (2011). https://doi.org/10.1586/epr.11.2

Stier A, Sackmann E. Spin labels as enzyme substrates Heterogeneous lipid distribution in liver microsomal membranes. Biochim. Biophys. Acta 311, 400–408 (1973). https://doi.org/10.1016/0005-2736(73)90320-9

Tanner W, Malinsky J, Opekarová M. In plant and animal cells, detergent-resistant membranes do not define functional membrane rafts. The Plant Cell 23, 1191–1193 (2011). https://doi.org/10.1105/tpc.111.086249

UniProt Consortium. UniProt: the universal protein knowledgebase. Nucleic Acids Res. 46, 2699 (2018). https://doi.org/10.1093/nar/gky092

Verma DK, Gupta D, Lal SK. Host Lipid Rafts Play a Major Role in Binding and Endocytosis of Influenza A Virus. Viruses 10, 650–661 (2018). https://doi.org/10.3390/v10110650

Wei X, She G, Wu T, Xue C, Cao Y. PEDV enters cells through clathrin -, caveolae -, and lipid raft - mediated endocytosis and traffics via the endo -/ lysosome pathway. Vet. Res. 51:10 (2020). https://doi.org/10.1186/s13567-020-0739-7

Westermann M, Leutbecher H, Meyer HW. Membrane structure of caveolae and isolated caveolin-rich vesicles. Histochemistry 111, 71–81 (1999). https://doi.org/10.1007/s004180050335

Xie Y, Zheng Y, Li H, Luo X, He Z, Cao S, Shi Y, Zhao Q, Xue Y, Zuo Z, Ren J. GPS-Lipid: a robust tool for the prediction of multiple lipid modification sites. Sci. Rep. 6, 28249 (2016). https://doi.org/10.1038/srep28249

Yang S-T, Kreutzberger AJB, Kiessling V, Ganser-Pornillos BK, White JM, Tamm LK. HIV virions sense plasma membrane heterogeneity for cell entry. Sci. Adv. 3, e1700338 (2017). https://doi.org/10.1126/sciadv.1700338

Yuan Z, Zhang F, Davis MJ, Bodén M, Teasdale RD. Predicting the solvent accessibility of transmembrane residues from protein sequence. J. Proteome Res. 5, 1063–1070 (2006). https://doi.org/10.1021/pr050397b

Yurtsever D, Lorent JH. Structural Modifications Controlling Membrane Raft Partitioning and Curvature in Human and Viral Proteins. J. Phys. Chem. B 124, 7574–7585 (2020). https://doi.org/10.1021/acs.jpcb.0c03435

