## Supplemental figures and tables for "ProRafts: A machine-learning predictor for *raftophilicity*, the protein affinity for biomembrane rafts"

#### **Supplementary Table**

Parameters used for Hyperparameter Tuning

| Parameter | Definition | List of Values | | | |
|---|---|---|---|---|---|
| | | **XGB - All** | **XGB - TM only** | **XGB - TM without TMD features** | **XGB - non-TM rafts** |
| min_child_weight | Minimum sum of instance weight (hessian) needed in a child. | [10, **20**, 35, 50] | [**10**, 15, 20, 25, 30, 35, 40] | [**10**, 15, 20, 25, 30, 35, 40] | [10, **20**, 35, 50] |
| gamma | Minimum loss reduction required to make a further partition on a leaf node of the tree. | [0, **2**, 4, 6, 8] | [0, **2**, 4, 6, 8] | [0, 2, **4**, 6, 8] | [0, 2, **4**, 6, 8] |
| max_depth | Maximum depth of a tree. | [4, 6, **8**] | [4, **6**, 8] | [2, **4**, 6, 8] | [4, **6**, 8] |
| n_estimators | Number of trees. | [35, 45, 50, 70, 80, 90, **100**] | [20, 30, 40, 45, 50, **60**, 110] | [20, 25, 30, 40, 45, 50, 60, 70, **75**] | [25, 40, 75, 85, 100, **130**] |
| learning_rate | Step size shrinkage used in update to prevent overfitting. | [0.01, 0.03, **0.05**] | [0.01, 0.03, **0.05**, 0.001] | [0.01, 0.03, **0.05**, 0.001] | [0.01, 0.03, **0.05**] |
| subsample | Subsample ratio of the training instances. | [**0.6**, 0.8, 1.0] | [***0.6****] | [***0.6****] | [0.6, ***0.7****, 0.8, 1.0] |
| colsample_bytree | Subsample ratio of columns when constructing each tree. | [0.6, ***0.7****, 0.8, 1.0] | [***0.6****] | [***0.6****] | [0.6, ***0.7****, 0.8, 1.0] |

**Parameter ranges for all four predictors used for Hyperparameter Tuning.** Table contains all the values used for hyperparameter tuning. The values in bold indicate the final parameter values chosen. The values marked with a star (*) were not among the initial parameters for hyperparameter tuning; these values were added during manual testing. The remaining parameters are the default values from XGBoost version 0.80.

### Supplementary Figure 1

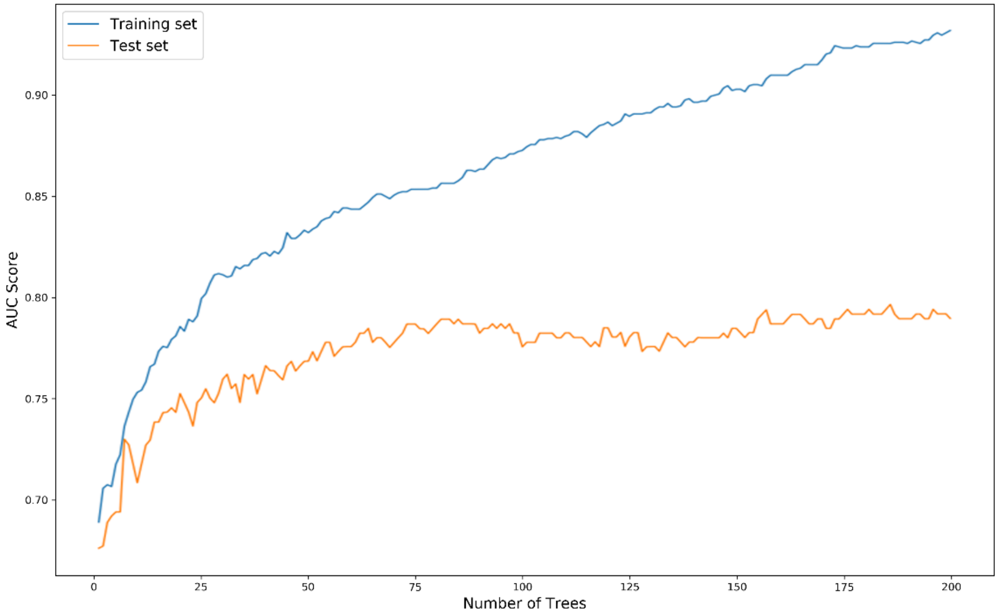


**Performance of the model by number of trees used.** Performance of the predictor, i.e., AUC score at the first step of the hyperparameter tuning, as a function of number of trees used (n_estimators) from 1 to 200. The performance of the model improves up to 100 trees; however, above this value, the AUC of the test dataset levels off, which indicates overfitting.

#### Supplementary Figure 2

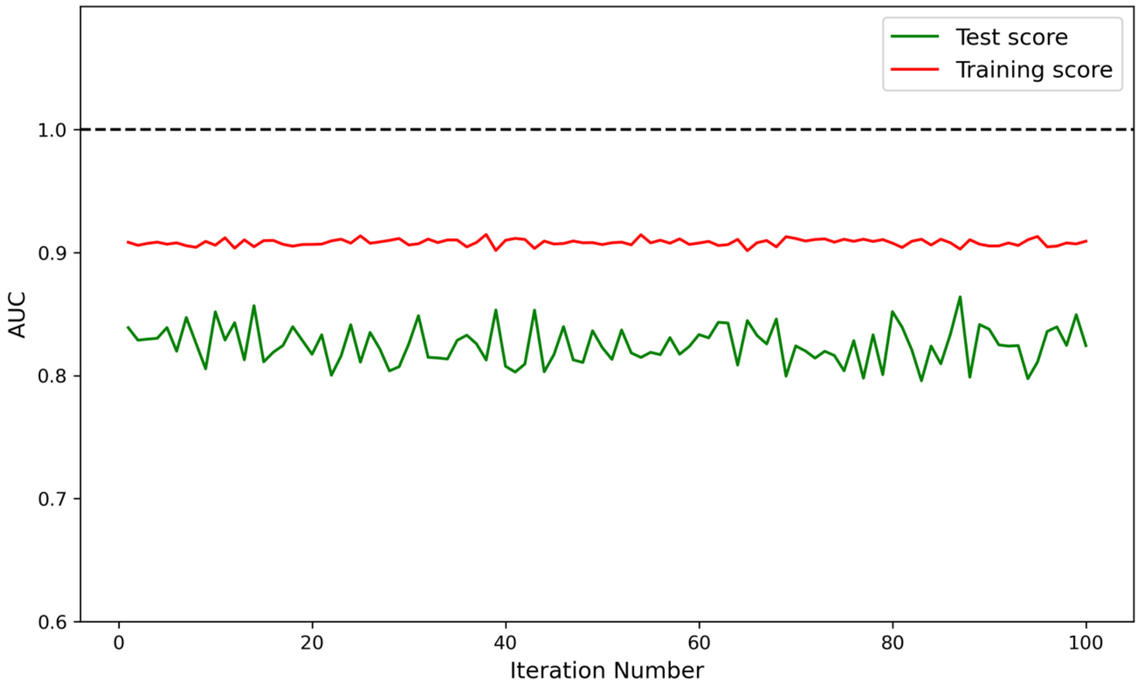


**Repeated trainings to test the effect of stochasticity on training** Due to the intrinsic stochasticity of the method and the small size of the dataset, prediction results show some variance. To understand better this variance, we trained the predictor 100 times, each time creating a new train-test split from the complete-set. The results are slightly variable though the prediction performance on the test set remains above 0.8 AUC.

### Supplementary Figure 3

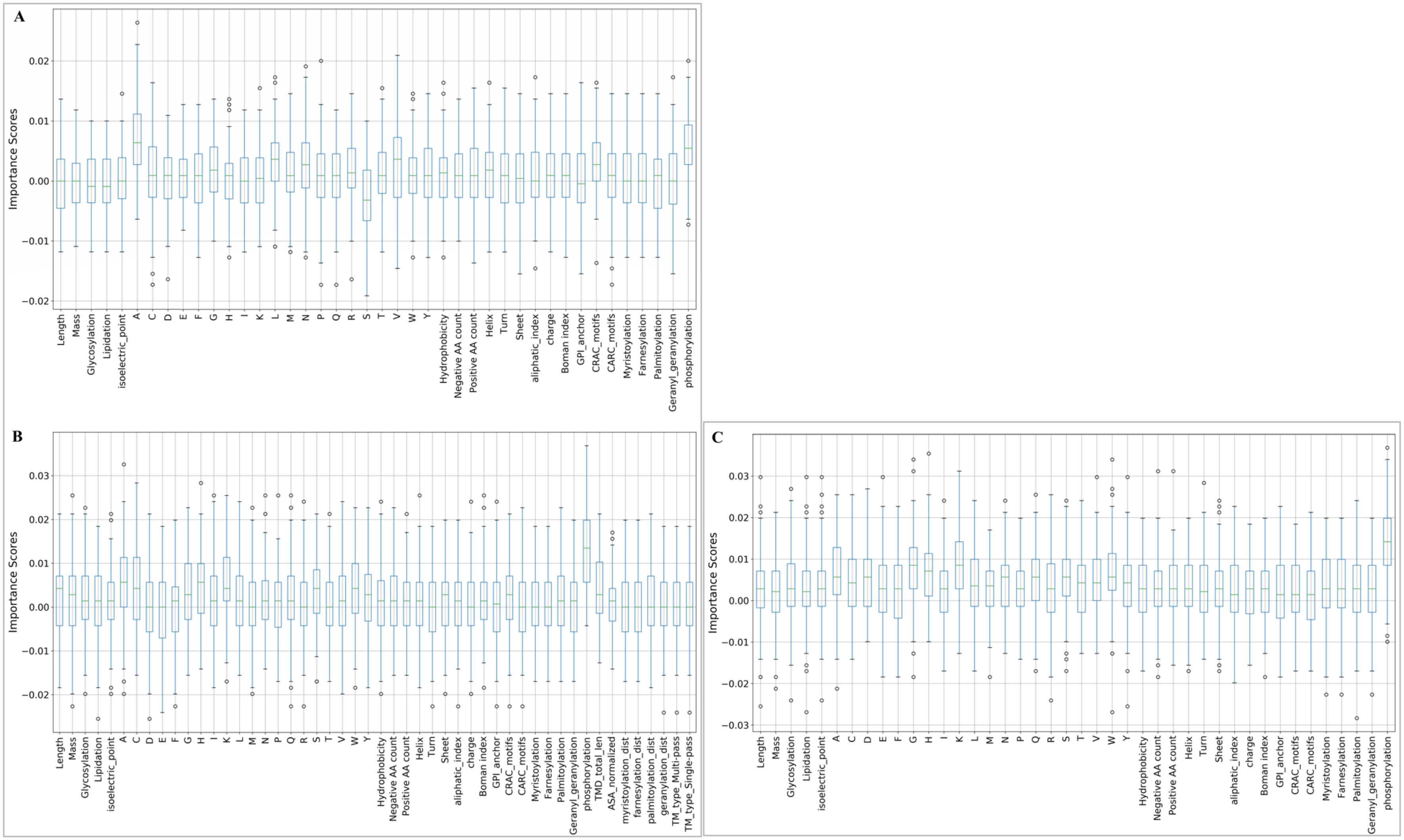


**Feature importance for the models trained on the sub-datasets.** Contribution of each feature to the prediction performance, obtained by training 100 times**. A,** training with non-TM raft proteins. **B,** training with TM only dataset. **C,** training with TM only dataset without specific TMD features. Each predictor make use of different sets of features; however, phosphorylation is the major contributor for all of them.
